# Correlated evolution of parental care with dichromatism, colour, and patterns in anurans

**DOI:** 10.1101/2021.04.11.439298

**Authors:** K S Seshadri, Maria Thaker

## Abstract

Parental care is widespread among vertebrates, with clear fitness benefits. Caring parents however incur costs that include higher predation risk. Anurans have among the most diverse forms of parental care, and we test whether the occurrence of care is associated with morphology that minimizes predation risk. We first examine whether parental care co-occurs with sexual dichromatism, testing the hypothesis that when one sex is conspicuous, the other is cryptically patterned and cares for the young. From our phylogenetic comparative analyses of 988 anurans distributed globally, we find that parental care is less likely to co-occur with dichromatism, irrespective of the caring sex. We then examine whether colour gradients and patterns that enhance crypticity are associated with the occurrence of parental care. We found that species with male-only care were more likely to have Bars-Bands, but contrary to our expectation, other colours (Green-Brown, Red-Blue-Black, Yellow) and patterns (Plain, Spots, Mottled-Patches) were not associated with caregiving behaviours. The lack of strong correlations between dorsal morphology and parental care suggests that crypticity is not the dominant strategy to minimise predation risk for care-giving anurans, and that the evolution of body colour and parental care are driven by independent selection pressures.

## Introduction

Understanding the patterns and processes underlying the evolution of parental care is central to evolutionary biology. Parental care in animals is an incredibly diverse trait, varying widely across taxa in both complexity and duration (Clutton-Brock 1991; Balshine 2012). Typically defined to include any post-zygotic investment by either parent (Trivers 1972), parental care includes both behavioural and non-behavioural traits that enhance offspring survivorship (Clutton-Brock 1991; Smiseth et al. 2012; Lindstedt et al. 2019). Currently, 11 forms of parental care are recognized across the animal kingdom and these range in complexity from provisioning of gametes and viviparity to protecting and feeding mature offspring (Smiseth et al. 2012). Inevitably, parental care is costly to the parent and theory predicts that caregiving behaviours evolve under circumstances when the benefits outweigh the costs (Clutton-Brock 1991; Alonso-Alvarez and Velando 2012; Royle et al. 2012; Klug and Bonsall 2014). Among the various costs, caregiving parents incur greater energetic expenditure and may undergo physiological stress (Fowler and Williams 2017). Parents also experience reduced resource acquisition opportunities, which can further reduce their allocation to other fitness-enhancing traits, including additional mating opportunities (Clutton-Brock 1991; Alonso-Alvarez and Velando 2012). Caregiving behaviours also exert direct mortality costs arising from the risks of predation not only to offspring but also to the adults themselves (Owens and Bennett 1994). In birds, specific caregiving behaviours such as feeding or brood defence are associated with higher predation risk compared to nest building or incubation (Owens and Bennett 1997). Depending on whether parents or offspring are at greater risk, caregiving parents can offset predation risk by avoiding detection or by actively engaging in antipredator strategies or both (Caro and Allen 2017).

One strategy to avoid detection and evade predation risk is through camouflage and the evolution of body colours and patterns that minimise predation risk is widespread in animals (Reviewed in Stevens and Merilaita 2009). Effective camouflage can be achieved in multiple ways: expression of colours and patterns that closely match the environment, achieving crypsis; expression of markings that create the appearance of false edges and boundaries, disrupting the outline and shape of the individual; development of body forms for masquerade; and integration of movement strategies to achieve the effects of motion dazzle or motion camouflage (Stevens and Merilaita 2011). A notable exception to morphology that minimizes predator detection is aposematic colouration (Mappes et al. 2005; Caro and Allen 2017).

Minimising detection with camouflage, however, is ineffective when an organism requires conspicuous body colour or patterns to communicate. Body colour patterns are often under strong sexual selection (Darwin 1871; Endler and Mappes 2017) and several vertebrates use visually striking colours as signals for both inter- and intraspecific communication (Bradbury and Vehrencamp 1998). In many taxa, such conspicuous body colours and patterns increase predation risk (Stuart-Fox et al. 2003), resulting in a trade-off between being prominent to attract mates, and being camouflaged to avoid predation (Endler 1980). Many species mediate this trade-off between sexual and viability selection by delaying the expression of conspicuous colours, restricting the duration of visual displays, dynamically exposing signals only to conspecifics, engaging in dynamic conspicuous colouration, or by evolving sexual dichromatism (Bell et al. 2017). Of these, sexual dichromatism is the most widely found evolutionary solution to the trade-off, wherein only the sex that experiences greater sexual selection is conspicuous while the other remains cryptic (Andersson 1994). Differences in crypticity between the sexes may be an interesting strategy to minimise predation risk for the parent that engages in parental care.

Amphibians are an ideal model system to examine the evolutionary trade-offs and associations between parental care and conspicuousness. Amphibians are a species-rich group (Wake and Koo 2018) that are colourful (Rojas 2017), dichromatic (Bell et al. 2017), and show some of the most diverse forms of caregiving behaviours (Wells 2007). Nearly 30 types of parental care behaviours have been documented in approximately 20% of the extant 8200 species of amphibians (Crump 2015; Furness and Capellini 2019; Vági et al. 2019; Schulte et al. 2020) and yet, our understanding of parental care evolution in amphibians is still lagging compared to other vertebrates (Stahlschmidt 2011). Even within amphibians, few studies examine the parental care behaviour of urodeles and caecilians compared to anurans (Schulte et al. 2020). What is clear is that caregiving behaviours among anurans are phylogenetically widespread and converge between distantly related lineages (Vági et al. 2019). Uniparental care, with males or females alone caring, is evolutionarily stable compared to biparental care (Furness and Capellini 2019). Across anuran species, caregiving behaviours vary in duration, intensity, and complexity (Crump 1995; Wells 2007; Delia et al. 2017; Vági et al. 2019). Among the nine classified categories of parental care in anurans (see: Schulte et al. 2020), egg attendance, wherein adults defend, clean, or hydrate eggs is the simplest and most frequently observed form (Wells 2007; Furness and Capellini 2019; Schulte et al. 2020) whereas viviparity is complex, requiring specialised anatomical and physiological adaptations that are infrequently observed (Furness and Capellini 2019; Schulte et al. 2020). Studies have found that parental care behaviour in anurans is correlated with multiple ecological factors, including breeding pool size and terrestrial reproduction (Brown et al. 2008; Brown et al. 2010; Vági et al. 2019) that are strong indicators of the risks to adults and young by predators in both the aquatic and terrestrial habitats.

Adult anurans are prey to visually hunting predators such as birds, snakes, mammals, and invertebrates (see: Toledo 2005; Toledo et al. 2007; Wells 2007) and avoiding predation risk by being camouflaged may explain the prevalence of green, brown, and grey dorsal colours that effectively match natural substrates (Wells 2007; Rojas 2017). Increasing crypticity of these dorsal colours is further aided by patterns, such as mottling, stripes, bars, or spots (Choi and Jang 2004; Nilsson Skold et al. 2013; Kang et al. 2016) and even metachrosis (Duellman and Trueb 1994). Several anuran families also effectively use aposematism to advertise unpalatability and avoid predation (Wells 2007; Rojas 2017). The adaptive significance of background matching by camouflage as well as aposematism to overcome predation is well supported at multiple scales (e.g., Summers and Clough 2001; Chiari et al. 2004; Choi and Jang 2004; Cooper et al. 2008; Hegna et al. 2012; Rojas et al. 2014; Taboada et al. 2020). Dichromatism could also be an effective strategy to overcome predation risk because the cryptically coloured sex could engage in caregiving behaviour while the other sex is conspicuous. Although evidence of the adaptive value of colours and patterns in shaping life-history traits is emerging, none, to the best of our knowledge, examine if costly life-history traits such as parental care are associated with dichromatism as well as specific dorsal colours, and patterns. Such a comparison is particularly relevant to understand how anurans ameliorate the trade-offs between viability selection and sexual selection.

Here, we examine whether the occurrence and type of parental care are evolutionarily correlated with dichromatism, and specific dorsal colours, and patterns in anurans. We first test for the co-occurrence of parental care and dichromatism, to examine whether dichromatism is more common in species where care is given by only males, only females, or both sexes. We also explore whether dichromatism is correlated with the developmental stage that receives care and the extent of protection. We then examine if parental care co-occurs with any of the three dorsal colour gradients and four pattern types. Given the relatively higher risks of conspicuous colouration, we hypothesized that caregiving adults are likely to be cryptically coloured and that specific patterns that enhance camouflage are more likely to be associated with caregiving behaviour. Using a large-scale phylogenetic comparison that links parental care to dichromatism, dorsal colour and pattern, we bring together seemingly independent traits to better understand the evolution of parental care and the potential trade-offs of sexual and viability selection that influence body colour morphology.

## Materials and Methods

Information on parental care in anurans was first extracted from data available in Vági et al. (2019), where we retained the following information: whether parental care is present or absent in each species; the type of care as one of 5 possible categories (no care, male-only care, female-only care, either parent, and biparental care); the developmental stage when care is provided as one of 4 possible categories (no care, egg care, tadpole care, and juvenile care); and extent of protection as one of 6 categories (no care, eggs in a nest without an attending parent, egg attendance by the parent, transport by carrying offspring on the body, transport by carrying offspring in the body, and viviparity). Because ‘care by either parent’ was rare (seven species only), we merged this category with ‘biparental care’ for analysis. The presence and form of dichromatism were listed from Bell and Zamudio (2012) and Bell et al. (2017). We augmented this list with information from the primary literature, field guides, and online databases (see electronic supplementary material, table S1). A species was classified as dichromatic if any part of the body was differently coloured compared to the other sex. We also recorded the specific body part that was dichromatic (e.g., vocal sac) if such information were available. We retained the classification of dichromatism made by Bell and Zamudio (2012), wherein ‘ontogenetic dichromatism’ between sexes is considered when males and females develop different colours at sexual maturity, and ‘dynamic dichromatism’ is when the male or female actively changes colour for a short duration at or after sexual maturity. We also recorded which sex (male or females) was dichromatic.

For dorsal colour and pattern information, one of us (SKS) examined colour photographs, illustrations, and referred to detailed descriptions on public repositories (see electronic supplementary material, table S1) of adult anurans. Because photos are taken under different lighting conditions, hues can vary and thus, photographs were used mainly to validate the description. The species description in the literature was our primary source for scoring colour categories. We classified the dorsal colour of males and females separately according to the following gradients: ‘Green-Brown’ which included hues of green or brown; ‘Red-Blue-Black’ which included hues of red or black or blue, and ‘Yellow’ which included all hues of yellow (figure 1). We combined red, blue and black into a single category because the independent occurrence of these colours was rare as they were often found in combination. We scored the pattern type into four categories *viz*., ‘Plain’: dorsum that lacks any pattern; ‘Bars-Bands’: dorsum with thin bars or wide bands; ‘Spots’: dorsum that is interspersed with distinct speckles or larger spots and, ‘Mottled-Patches’: dorsum with blotches or indistinct shaped patches (figure 1). The colour and pattern of polymorphic species (< 1 % of all species in our database) were classified from the type specimen, described in the original species descriptions. These colour gradients enabled us to categorize species into broad colour schemes that were not directly quantifiable from specimens or photographs or illustrations as others have done (e.g., Dale et al. 2015). The four pattern categories are reliably distinguishable by observers and have been used successfully to address similar questions in reptiles (see: Allen et al. 2020). After removing species for which reliable information on dichromatism, colour, and pattern was unavailable, our final dataset comprised 988 species belonging to 45 families. Species names follow Frost (2020).

**Figure 1.**
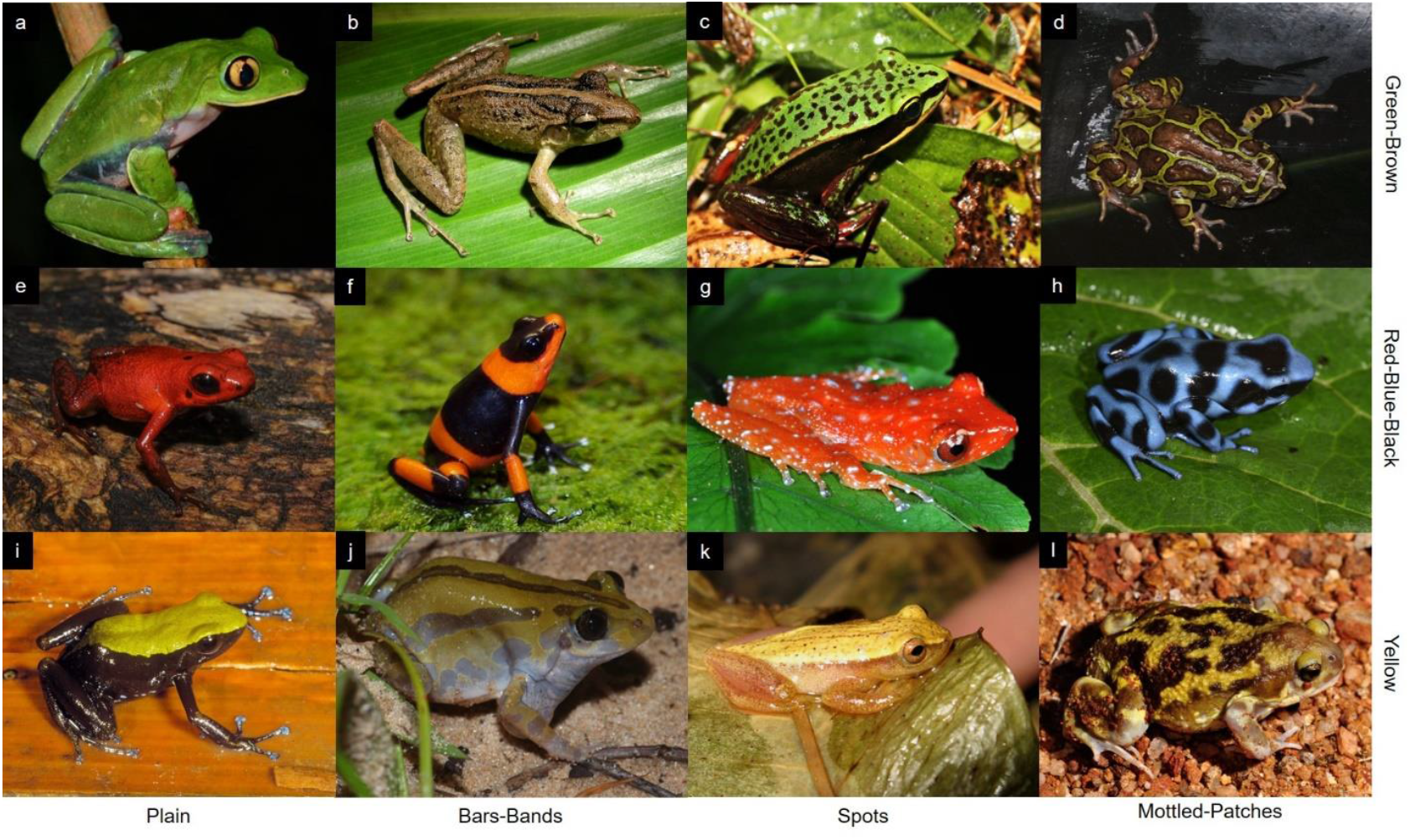
Combinations of the three colours (rows), and four dorsal patterns (columns) categorised for anurans in our dataset: a. Green-Brown with plain dorsum in *Agalychnis annae* (Adam W. Bland); b. Green-Brown with Bars-Bands in *Craugastor fitzingeri* (Brian Kubicki); c. Green-Brown with Spots in *Lithobates vibicarius* (David A. Rodríguez); d. Green-Brown with Mottled-Patches in *Scaphiophryne marmorata* (Peter Janzen); e. Red-Blue-Black with Plain dorsum in *Andinobates opisthomelas* (Daniel Vásquez-Restrepo); f. Red-Blue-Black with Bars-Bands in *Oophaga lehmanni* (Arachnokulture); g. Red-Blue-Black with Spots in *Nyxtixalus pictus* (Seshadri KS); h. Red-Blue-Black with Mottled-Patches in *Dendrobates auratus* (Peter Janzen); i. Yellow with Plain dorsum in *Mantella laevigata* (Miguel Vences); j. Yellow with Bars-Bands in *Kassina senegalensis* (Alberto Sanchez-Vialas); k. Yellow with Spots in *Dendropsophus sanborni* (Raul Maneyro) and; l. Yellow with Mottled-Patches in *Uperodon systoma* (Seshadri KS). All images except g & l are sourced from CalPhotos with permission from respective owners (credits in parentheses).

### Comparative analyses

To examine the evolutionary association of parental care and body colouration, we modified the consensus tree provided by Jetz and Pyron (2018) to only include the 988 species in our database. Data on dichromatism and parental care were plotted on a phylogenetic tree with branch lengths transformed by Graphen’s ‘ρ’ (rho) for illustrations (Grafen 1989). Data on parental care, type of care, dichromatism, colour, and pattern were binary, and therefore we used a phylogenetic generalized linear mixed model for binary data using the function BinaryPGLMM (Paradis et al. 2004). We ran six phylogenetically corrected comparative models to test for correlations. The models returned estimated regression coefficients, standard errors, and strength of the phylogenetic signal (s^2^) based on a variance-covariance matrix (Ives and Helmus 2011). The occurrence of parental care (care or no care) and type of care (no care, male-only, female-only, and biparental care) were treated as dependent variables. Dichromatism (present or absent), dorsal colour (Green-Brown or Red-Blue-Black or Yellow) and, pattern (Plain, Bars-Bands, Spots, Mottled-Patches) with the latter two variables for males and females separately, were treated as independent variables. All analyses were performed using RStudio v 1.3 (RStudio Team 2020) using the following packages ‘caper’ (Orme et al. 2013), ‘ape’ (Paradis et al. 2004), ‘geiger’ (Pennell et al. 2014), ‘phytools’ (Revell 2012), ‘ggtree’ (Yu et al. 2017).

## Results

### Overview of Parental care, Dichromatism, Body Colour, and Pattern in anurans

Parental care was present in 375 species (out of 988 species) wherein male-only care was most frequent (n = 204) followed by female only (n = 106), and biparental care (n = 65). Dichromatism was observed in 221 species and the ontogenetic form (n = 131) was more frequent than dynamic dichromatism (n = 90). Overall, Green-Brown was the most common dorsal colour gradient (male = 720, female = 720) followed by Red-Blue-Black (male = 146, female = 150), and Yellow (male = 122, female = 118). Unlike dorsal colours, frequency of different dorsal pattern types was more even in these anurans, with Mottled-Patches being the most common (male = 303, female = 303) followed by Spots (male = 243, female = 250), Bars-Bands (male = 225, female = 218), and Plain (male = 217, female = 217).

### The association of parental care with dichromatism

Dichromatism was widespread across the phylogeny (figure 2a, electronic supplementary material, figure S1) with a strong phylogenetic signal (s^2^ = 20.62, p < 0.005). The occurrence of parental care was more strongly associated with non-dichromatic species than dichromatic species (figure 2b; table 1, model 1). Similarly, types of care (no care, male-only, female-only, biparental) were distributed widely across the phylogeny (figure 2c) with a moderately strong phylogenetic signal (s^2^ = 3.3, p < 0.005). All three types of parental care were negatively associated with the presence of dichromatism, but more specifically, this negative association was statistically significant with male-only and female-only care (figure 2d; table 1, model 2). When we examine the developmental stages that receive care, we find that care during the egg stage is the most common in both dichromatic and non-dichromatic species, followed by care at the tadpole and juvenile stages (figure 3a). Similarly, both dichromatic and non-dichromatic species that do show parental care mainly attend to eggs or transport offspring on their back (figure 3b)

**Figure 2.**
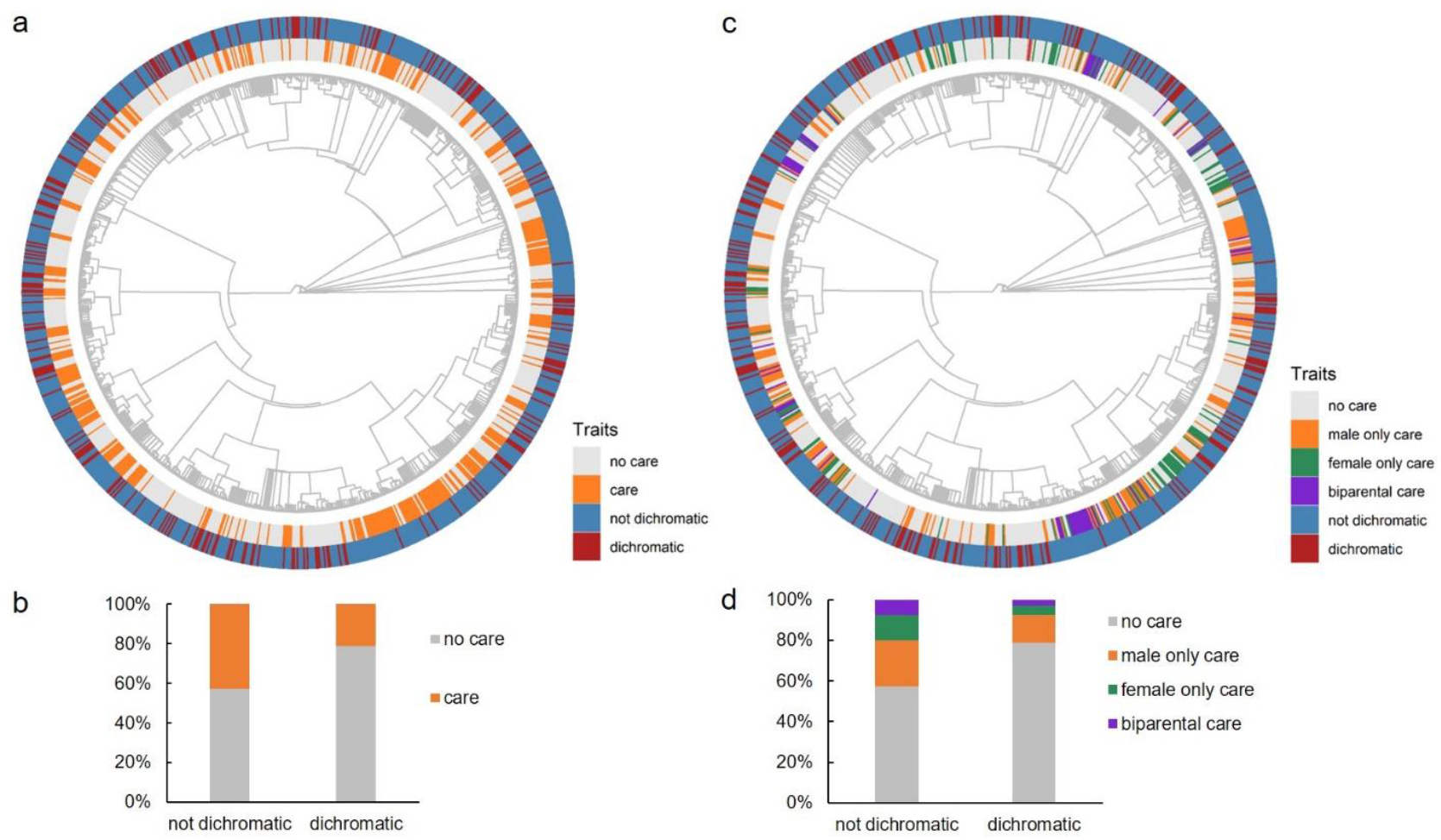
The association between dichromatism and parental care in anurans (N = 988 species). (a) Phylogenetic distribution and (b) co-occurrence of dichromatism (presence or absence) with parental care (care or no care). (c) Phylogenetic distribution and (d) co-occurrence of dichromatism (presence or absence) with different types of parental care (no care, male-only care, female-only care, biparental care). (b) and (d) report % of species.

**Table 1.** Evolutionary associations between the occurrence and type of parental care and the presence of dichromatism, and gradients of dorsal colour and pattern. Shown are outputs of binaryPGLMM regressions with the dependent and explanatory variables listed according to the models tested. P < 0.05 are indicated by an ‘*’. All models tested data from 988 species. N is the number of species having the condition listed as a dependent variable, e.g., parental care was present in 375 species.

| Model no | Dependent | Explanatory | Estimate ± SE | Z | P | Phylogenetic signal | Pr | N |
| --- | --- | --- | --- | --- | --- | --- | --- | --- |
| 1 | Parental care | Not dichromatic (Intercept) | $-0.5 \pm 1.6$ | -0.30 | 0.76 | 20.62 | 1.00E-41 | 375 |
| | | Dichromatic | $-0.71 \pm 0.3$ | -2.15 | 0.03* | | | |
| 2 | Dichromatism | No care (Intercept) | $-1.68 \pm 0.7$ | -2.29 | 0.02* | 3.319 | 2.30E-38 | 221 |
| | | Male only care | $-0.83 \pm 0.3$ | -2.51 | 0.01* | | | |
| | | Female only care | $-0.96 \pm 0.4$ | -2.29 | 0.02* | | | |
| | | Biparental care | $-0.98 \pm 0.5$ | -1.86 | 0.06 | | | |
| 3 | Male only care | Plain (Intercept) | $-2.2 \pm 1.6$ | -1.39 | 0.16 | 1.85E+01 | 5.25E-57 | 204 |
| | | Bars-Bands on males | $0.9 \pm 0.4$ | 2.33 | 0.02* | | | |
| | | Spots on males | $0 \pm 0.4$ | 0.1 | 0.92 | | | |
| | | Mottled-Patches on males | $0.4 \pm 0.4$ | 1.19 | 0.24 | | | |
| 4 | Male only care | Green-Brown (Intercept) | $-1.8 \pm 1.5$ | -1.13 | 0.26 | 18.63 | 2.94E-60 | 204 |
| | | Red-Blue-Black on males | $-0.3 \pm 0.3$ | -0.87 | 0.39 | | | |
| | | Yellow on males | $-0.2 \pm 0.3$ | -0.54 | 0.59 | | | |
| 5 | Female only care | Plain (Intercept) | $-0.3 \pm 1.5$ | -1.94 | 0.05 | 15.01 | 1.35E-67 | 106 |
| | | Bars-Bands on females | $-0.4 \pm 0.5$ | -0.96 | 0.34 | | | |
| | | Spots on females | $-0.2 \pm 0.4$ | -0.35 | 0.73 | | | |
| | | Mottled-Patches on females | $-0.3 \pm 0.4$ | -0.71 | 0.48 | | | |
| 6 | Female only care | Green-Brown (Intercept) | $-3.3 \pm 1.5$ | -2.14 | 0.03* | 15.19 | 2.42E-67 | 106 |
| | | Red-Blue-Black on females | $0.2 \pm 0.4$ | 0.5 | 0.62 | | | |
| | | Yellow on females | $0.2 \pm 0.5$ | 0.4 | 0.69 | | | |

**Figure 3.**
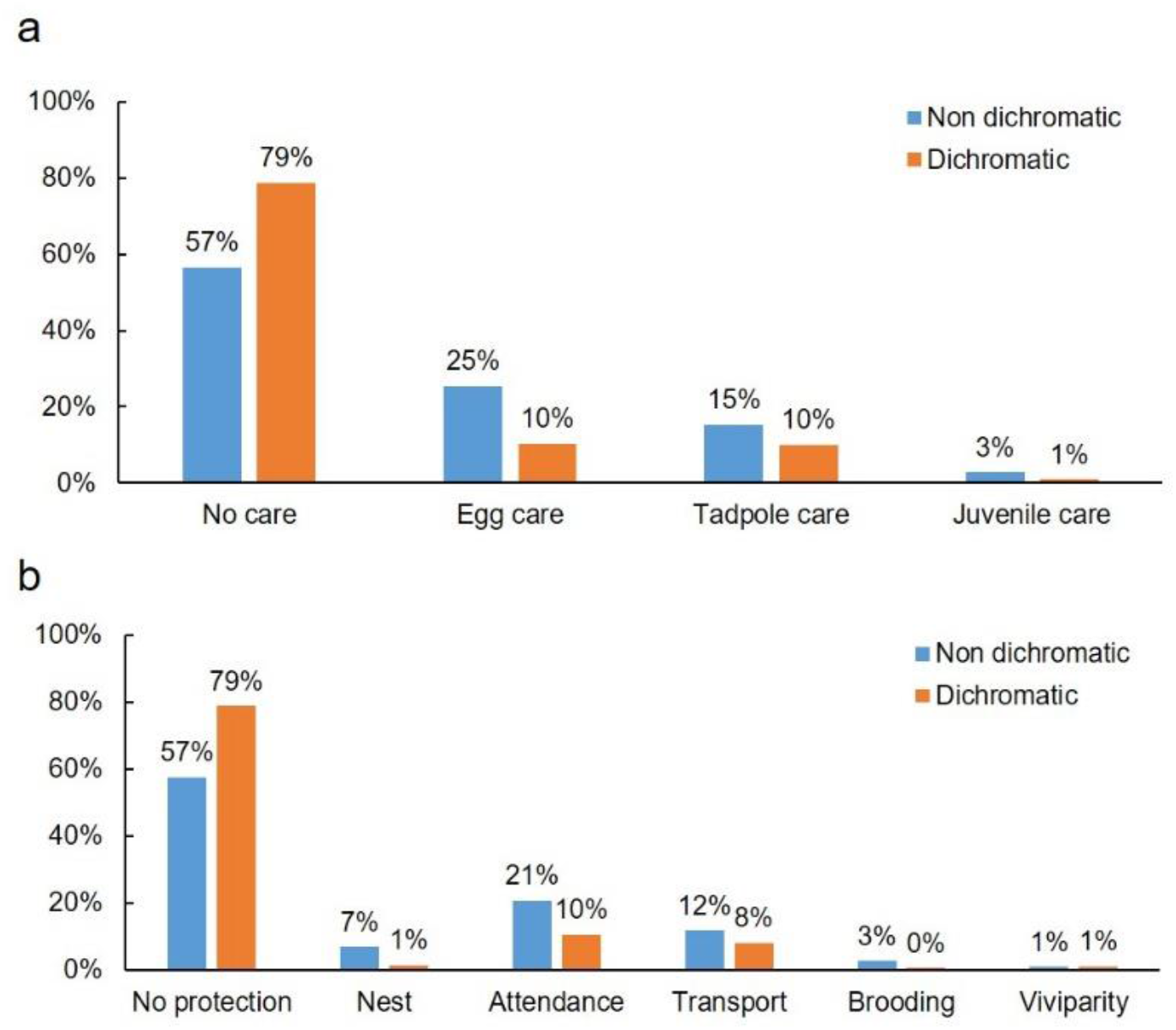
Forms of parental care can vary depending on the (a) developmental stage of the offspring: no care, eggs, tadpoles or juveniles, and (b) the extent of protection provided: no care, eggs in a nest but without attendance by parents, egg attendance, transport on the back, transport inside the body or brooding, and viviparity. Shown are relative percentages of these caregiving forms within dichromatic (n = 221) and non-dichromatic (n = 767) species.

### The association between type of care with dorsal colours and patterns

Irrespective of whether species are dichromatic or not, those that exhibit parental care varied widely in body colours (figure 4a, b) and patterns (figure 4c, d), with no strong correlations. By contrast, species with male-only care had a strong phylogenetic signal and was significantly correlated with the presence of Bars-Bands on those males (table 1, model 3). We found no significant correlations between male-only care and any of the three colour gradients (table 1, model 4). Female-only care was also not significantly correlated with any of the three colours or four patterns (table 1, model 5,6).

**Figure 4.**
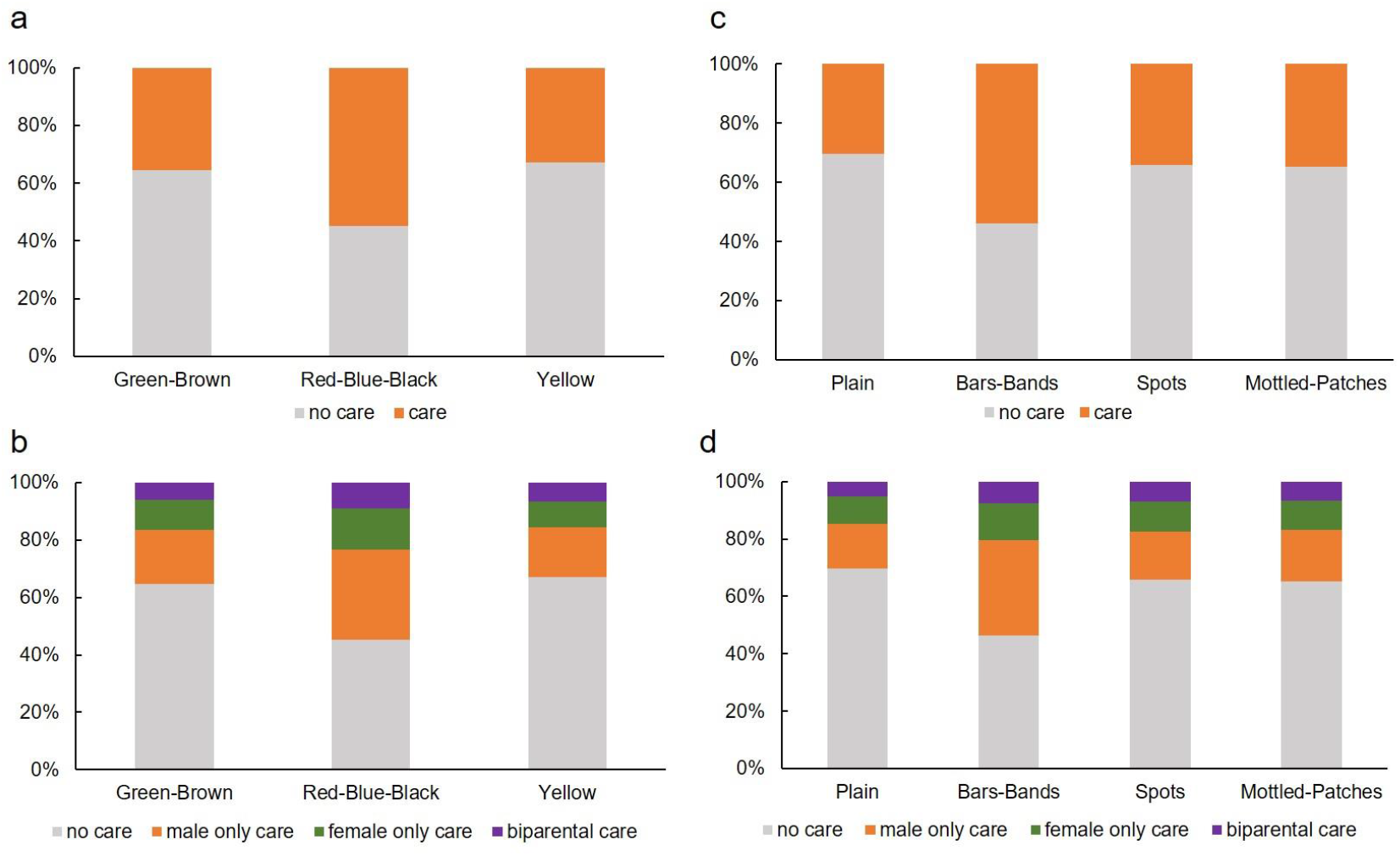
Parental care (no care, care) and type of parental care (no care, male-only care, female-only care, and biparental care) are weakly correlated with (a, b) dorsal colour and (c, d) dorsal pattern, except for the significant association of male-only care with Bars-Bands. Shown are relative percentages of each combination of traits of males from 988 species.

## Discussion

Our understanding of the evolution of anuran parental care has largely been focussed on determining costs and benefits (Alonso-Alvarez and Velando 2012) or the macro-evolutionary comparisons with other ecological factors, such as terrestriality (Vági et al. 2019). Here, we aimed to test if parental care behaviours were correlated with morphological traits under both viability and sexual selection. Our large-scale phylogenetic analysis revealed that dichromatism is extremely variable among anurans but overall, a greater number of non-dichromatic species care for their offspring than dichromatic species. We also found that species with male-only care were more likely to have Bars-Bands on their dorsum, irrespective of dorsal colour. No correlations between female-only care and biparental care were found with dorsal colour and pattern. Differences in the associated evolution of some morphological traits with parental care by males and females suggests interesting sex-specific differences in the trade-offs between viability and sexual selection. Taken together, these findings suggest that the occurrence of parental care among anurans is more strongly associated with non-dichromatic species, but the colours and patterns of caregiving species are not tightly linked to the presence of care overall.

### Parental care and Dichromatism

Our results show that dichromatic species are less likely to care for the offspring compared to non-dichromatic species, and the pattern is consistent irrespective of the caregiving sex. Dichromatism has been recorded in ∼2% of extant anurans (Portik et al. 2019), but this is likely to be an underestimate, owing to the ephemeral nature of dynamic dichromatism (Rojas 2017) and the general lack of studies (Rojas 2017; Wake and Koo 2018). Typically, dichromatism involves a change to a bright, conspicuous colour on the body either with age or sexual maturity or during the breeding season. Dichromatism among anurans is known to be evolutionarily labile and has been lost and gained multiple times over their evolutionary history (Bell et al. 2017). However, only two studies have so far examined the macroevolutionary patterns of dichromatism in anurans: one highlights the presence of dichromatism among anuran lineages and contextualizes existing knowledge (Bell and Zamudio 2012), while the other finds that the evolution of dynamic dichromatism and explosive breeding are correlated (Wells 1977; Bell et al. 2017). The evolution of dichromatism appears to have been preceded by breeding in aggregations (Bell et al. 2017), therefore enabling male-male recognition (Sztatecsny et al. 2012; Kindermann and Hero 2016) as well as female assessment of male quality (Gomez et al. 2009). Anurans that form aggregations may not engage in parental care behaviours such as egg attendance or froglet transport that inherently require prolonged association with immobile offspring (Wells 2007). Some dichromatic anurans may be capable of prioritizing both sexual and viability selection using an ingenious solution of restricting the conspicuousness to a small portion of the body (see electronic supplementary material, table S1 for references) such as feet (Forelimbs turn orange in males of *Pyxicephalus adspersus*), throat (Males of *Phrynoidis asper* have black throat), or the ventral surface (Males of *Hyloxalus delatorreae* have a spotted underside). We suspect that this strategy of restricted dichromatism may reduce the risk of predation further because the conspicuous signal is directed specifically to a receiver and may only be exposed in close encounters, thus avoiding the attention of unintended recipients. Overall, the negative correlation between dichromatism and parental care seems to suggest that parental care is incompatible with the sexually selected trait of dichromatism among anurans. Given that parental care and dichromatism require complex morphological, physiological, and behavioural adaptations that affect their relative lability (Furness and Capellini 2019; Taboada et al. 2020), analysing the evolutionary transitions to each state would enable us to draw interesting inferences on the sequence of evolution of these two traits.

### Parental care, colour, and pattern

The presence of Green-Brown colours is thought to render advantages for camouflage to anurans (Wells 1977; Taboada et al. 2020). Contrary to our expectation, we did not find caregiving species to be those that are also green or brown. None of the three colour gradients were significantly associated with the occurrence of care, irrespective of the caregiving sex. The absence of an association between parental care and dorsal colour may reflect the many other functions of body colouration. Red colour, for instance, is a ubiquitous warning colour across different taxonomic groups (Mappes et al. 2005) and is an effective deterrent of predation as in the case of the Strawberry poison dart frog, *Oophaga pumilo*, where the green morph is more likely to be attacked than the red morph (Hegna et al. 2012). Although bright or high-contrast colours have generally been associated with aposematism, some anurans such as *Dendrobates tinctorius* use yellow and black patterns in a dual role as disruptive camouflage as well as aposematism (Barnett et al. 2018). Furthermore, several palatable species use aposematic colouration to mimic unpalatability (e.g. Uropeltid snakes Cyriac and Kodandaramaiah 2019). What is particularly informative of our results is that parental care was not associated with species that were either cryptically coloured or conspicuously coloured. Because the effectiveness of colours in reducing predation risk depends on the visual acuity of predators and the visual complexity of habitats, our evolutionary hypotheses about patterns of these morphological traits and parental care may be better understood by including current ecological conditions.

Of the four pattern categories we classified in anurans, only the presence of Bars-Bands had a significant positive association with male-only care. In contrast, the association of this pattern with female-only care was weak and negative. None of the other three pattern types, *viz*., Plain, Mottled-Patches, or Spots had a significant association with the sex of the caregiver. We suspect that the differences in the association between male-only and female-only care may be an artefact of fewer species having female-only care compared to male-only care. However, the fact that at least in males, Bars-Bands could be an added advantage for the care-giving parent, is indicative of a sex-specific trade-off between viability selection and sexual selection. In allied taxa such as geckos and snakes, bars-bands or zig-zag patterns on the dorsum are known to enhance survival, irrespective of body colour (Wuster et al. 2004; Niskanen and Mappes 2005). Among geckos, the presence of bars and bands on the dorsum is thought to enable background matching (Allen et al. 2020). Male anurans that occupy linear habitats such as grass or tree bark may also benefit from background matching. Mottled patterns and spots on the dorsum would likely match backgrounds such as mud or leaf litter (Rojas 2017), but the effectiveness of different patterns for background matching in anurans across habitats remain to be quantified. It also remains to be seen if males and females occupy different oviposition sites or microhabitats depending on their specific dorsal pattern. Our categorisation of colour and patterns into three and four categories respectively may not necessarily reflect the true extent of camouflage or conspicuousness as perceived by predators, especially since thermal conditions also modulate colour patterns (Wells 2007). We also recognise that our findings are based on colour and patterns perceived in the visible spectrum and there is mounting evidence of some anurans using the ultra-violet spectrum (UV) and being fluorescent (Taboada et al. 2017). Finally, because of the general lack of data, what is missing is information about the diurnal or nocturnal habit of anurans. Several conspicuously coloured anurans are diurnal but nocturnality may inherently enhance crypsis and reduce detection by diurnal predators. The adaptive values of colours and patterns will be influenced by whether these anurans are conspicuous and at risk to their predators when they are active and expressing parental care.

### Predation risk and aversive strategies while caring for offspring

Predation is a key driver for the evolution of parental care in amphibians (Wells 2007). There is strong evidence showing that parental care behaviours increase offspring survivorship (e.g., Townsend 1986; Bickford 2002; Bickford 2004; Poo and Bickford 2013; Delia et al. 2017; Seshadri and Bickford 2018; Delia et al. 2020; Schulte et al. 2020). However, little is known about the magnitude of predation risk on caregiving parents themselves. Many caregiving anuran species, such as *Feihyla hansenae*, actively defend their eggs from predators such as Katydids (Poo et al. 2016). Anurans can use a range of strategies to reduce predation risk: by abandoning risky oviposition sites (Chuang et al. 2017); evolving morphology and behaviours that enhance crypticity; and actively evade capture once detected (see: Toledo et al. 2011). Documenting the suite of predators and predation rates on different colours and patterns of caregiving adult anurans, as well as a careful evaluation of life-history traits such as the presence of toxicity and aposematism, may reveal further insights on combinations of anti-predator strategies that anurans can utilise. Experiments that evaluate the risk of predation across the three colours and four dorsal patterns among anurans will be particularly useful to understand if specific combinations of colour and pattern indeed enhance camouflage. From our phylogenetic analysis, the lack of strong correlations between the expression of parental care and specific dorsal colours (Green-Brown, Red-Blue-Black and, Yellow) and patterns (Plain, Spots, and Mottled-Patches) reflects an interesting disassociation between colour morphology and parental care. The evolution of care-giving behaviour does not appear to preclude the evolution of the myriad colour and patterns characteristic of anurans globally.

## Supporting information

electronic supplementary material, table S1

electronic supplementary material, figure S1

## Acknowledgements

We appreciate the insightful comments and discussions with Priya Iyer, Harish Prakash, and Vidisha M.K. on early versions of this manuscript.

