## Supplementary figures and images for "Correlated evolution of parental care with dichromatism, colour, and patterns in anurans"

### electronic supplementary material, figure S1

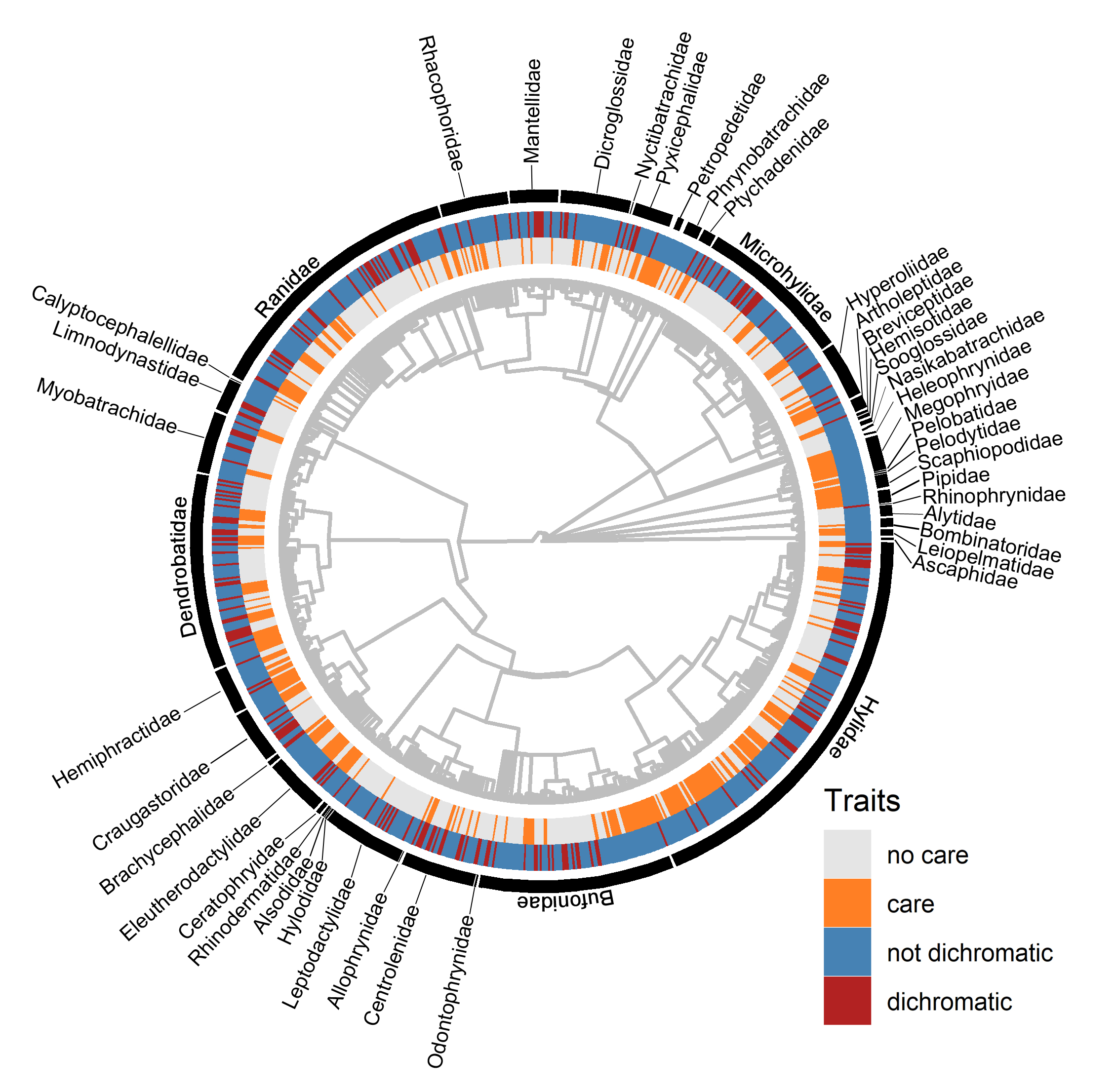
